# A new paradigm of intracrine free fatty acid receptor 4 signaling at lipid droplets

**DOI:** 10.1101/2023.07.28.550805

**Authors:** Emma Tripp, Shannon L. O’Brien, Gabrielle Smith, Adam Boufersaoui, Jennie Roberts, Jeremy Pike, Joao Correia, Tamara Miljus, Daniel A. Tennant, Brian D. Hudson, Graeme Milligan, Zachary Gerhart-Hines, Thue W. Schwartz, Davide Calebiro

**Affiliations:** Institute of Metabolism and Systems Research, University of Birmingham, Birmingham, United Kingdom; Centre of Membrane Proteins and Receptors (COMPARE), Universities of Nottingham and Birmingham, Birmingham, United Kingdom; Centre for Translational Pharmacology, School of Molecular Biosciences, College of Medical, Veterinary and Life Sciences, University of Glasgow, Glasgow, UK; Novo Nordisk Foundation Center for Basic Metabolic Research, University of Copenhagen, Copenhagen, Denmark

## Abstract

G protein-coupled receptors (GPCRs), once thought to be active exclusively at the plasma membrane, have been shown to signal from multiple intracellular membrane compartments, including endosomes and the Golgi. However, the potential occurrence and functional relevance of intracellular signaling for the emerging family of metabolite-sensing GPCRs is largely unknown. Here, we used live-cell imaging, bioluminescence resonance energy transfer (BRET) measurements, and functional readouts to investigate signal compartmentalization of the free fatty acid receptor 4 (FFA4), a prototypical metabolite-sensing GPCR that is activated by medium- and long-chain free fatty acids (FFAs). Unexpectedly, we show that FFA4 largely resides on intracellular membranes that are intimately associated with lipid droplets in adipocytes. Upon lipolysis induction, the released FFAs rapidly bind to and activate this intracellular pool of FFA4, leading to local G_i/o_ coupling and inhibition of cAMP production in the vicinity of lipid droplets. This provides a spatiotemporally confined negative feedback mechanism allowing individual lipid droplets to rapidly adjust their lipolysis rate. Our results reveal a novel ‘intracrine’ signaling modality by a prototypical metabolite-sensing GPCR and identify a new lipid-droplet-associated signaling hub implicated in the rapid regulation of lipid metabolism, with important implications for adipocyte physiology and pharmacology.

## INTRODUCTION

Metabolites function as essential energy sources and building blocks for biosynthetic pathways^1^. Classically, they were thought to modulate metabolic pathways primarily via directly regulating the activity of key biosynthetic enzymes. However, with the deorphanization of several G protein-coupled receptors (GPCRs) with previously unknown endogenous ligands, it has become apparent that metabolites can also function as bona fide agonists for a growing number of metabolite-sensing GPCRs that modulate cellular functions in a similar manner to receptors for hormones and neurotransmitters^1,2^.

In parallel, although originally believed to be active only at the plasma membrane, evidence accumulated over the past 15 years indicates that GPCRs can also signal via G proteins from intracellular sites such as early endosomes or the Golgi/*trans*-Golgi network to locally control cellular functions^3-8^. Given the emerging paradigm of GPCR signaling at intracellular sites and the compartmentalized nature of intracellular metabolism^9,10^, an intriguing hypothesis is that metabolite-sensing GPCRs might signal from intracellular membranes in order to locally regulate metabolic pathways. However, the potential occurrence and physiological relevance of metabolite-sensing GPCR signaling at intracellular sites remains to be clarified.

A prime example of a tightly regulated metabolic pathway is lipolysis, which is stimulated by hormones like adrenaline acting upon G_s_-coupled receptors present on the surface of adipocytes. The resulting increase in intracellular cyclic AMP (cAMP) levels and protein kinase A activation trigger the release of free fatty acids (FFAs) that are stored as triglycerides in lipid droplets. FFAs, in turn, have been shown to activate the free fatty acid receptor 4 (FFA4), also known as GPR120, a GPCR that is highly expressed in adipocytes^11- 14^ and is activated by medium- and long-chain FFAs^15^. Recent studies suggest that the FFA4 might play an important role in modulating adipocyte metabolism by forming part of a feedback loop that negatively modulates lipolysis^16,17^. However, the underlying molecular mechanisms are insufficiently understood.

Here, we use a combination of real-time bioluminescence resonance energy transfer (BRET) measurements, live-cell imaging, and functional read-outs to investigate the spatiotemporal organization of FFA4 signaling in adipocytes. Surprisingly, our results reveal that FFA4 resides and signals at intracellular sites that are intimately associated with lipid droplets, suggesting that it forms part of an ‘intracrine’ GPCR feedback mechanism capable of modulating lipolysis via locally inhibiting cAMP signaling. These results provide direct evidence for intracrine signaling by a metabolite-sensing GPCR and identify a novel lipid droplet-associated GPCR signaling hub implicated in the rapid regulation of lipid metabolism in adipocytes.

## RESULTSs

### FFA4 is primarily coupled to Gα_i/o_ in a simple cell model

FFA4 was originally thought to be predominantly coupled to G_q/11_, mainly due to the observation of Ca^2+^ increases upon pharmacological receptor activation^18,19^. However, recent studies have also suggested a possible coupling to Gα_i/o_^17,20,21^, which could provide a more direct mechanism for the inhibitory effects of FFA4 on lipolysis. To further investigate the coupling specificity of FFA4, we took advantage of mini-G probes, which consist of engineered GTPase domains of Gα subunits that rapidly translocate from the cytoplasm to active GPCRs upon receptor activation^22,23^. Since mini-G probes retain coupling specificity, they can be used to study GPCR coupling in living cells^23^. For this purpose, we performed real-time BRET measurements between FFA4 carrying NanoLuc luciferase (Nluc) at its C-terminus (FFA4-Nluc) and Venus-tagged mini-G probes (mGα_i_, mGα_o_, mGα_q_, mGα_s_, or mGα_12_) expressed in human embryonic kidney 293T (HEK293T) cells (Fig.1A). Stimulation with two selective synthetic FFA4 agonists, TUG-891 (Fig. 1B, C)^19^ and Compound A (CpdA) (Fig. 1C, Supplementary Fig. 1A)^24^, as well as the endogenous agonist α-linolenic acid (Fig. 1C, Supplementary Fig. 1A)^15^, caused a robust increase in BRET between the receptor and both mGα_i_ and mGα_o_. In contrast, only modest increases were observed with mGα_q_ and to a lesser extent the remaining mini-G probes (Fig. 1B, C, Supplementary Fig. 1A). These results indicate that FFA4 is strongly coupled to Gα_i/o_ in a simple cell model.

**Figure 1:**
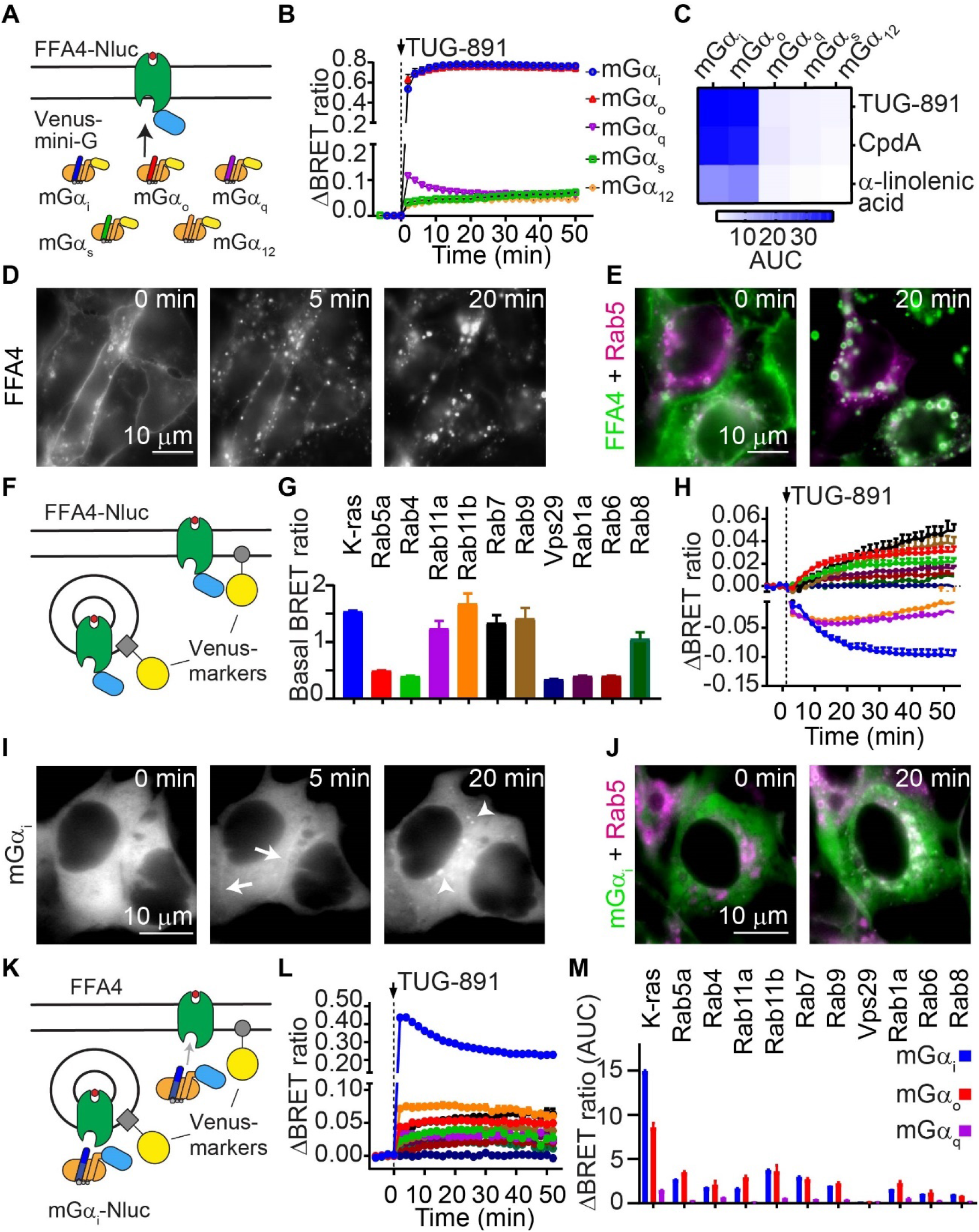
FFA4 rapidly traffics to and signals via Gα_i/o_ from intracellular compartments after agonist stimulation in a simple cell model. HEK293T cells were transfected with the indicated constructs and stimulated with 10 μM TUG-891. **(A)** Schematic of the BRET assay used to measure FFA4 coupling specificity with mini-G probes. **(B)** Agonist-dependent increases in Venus-tagged mini-G probe recruitment to FFA4-Nluc upon TUG-891 stimulation, monitored by real-time BRET. **(C)** Heat map comparing the AUC values of mini-G probe recruitment with TUG-891, Compound A (CpdA) or α-linolenic acid. **(D)** Effect of TUG-891 stimulation on FFA4 localization. Shown are selected frames from a representative HILO image sequence of cells transfected with FFA4-YFP. **(E)** Colocalization of FFA4-YFP with an early endosome marker (Rab5-mCherry) in cells stimulated with TUG-891. White color indicates colocalization. **(F)** Schematic illustrating the bystander BRET assay used to monitor FFA4 internalization and intracellular trafficking in real time. **(G)** Results of basal bystander BRET measurements between FFA4-Nluc and Venus-tagged subcellular markers. **(H)** Agonist-dependent changes in FFA4 localization at subcellular compartments, monitored by real-time bystander BRET between FFA4-Nluc and Venus-tagged subcellular markers. **(I)** Effect of TUG-891 stimulation on Venus-mGα_i_ translocation. Shown are selected frames from a representative HILO image sequence of cells co-transfected with FFA4 and Venus-mGα_i_. Arrows, plasma membrane. Arrowheads, endosomes. **(J)** Colocalization of Venus-mGα_i_ with an early endosome marker (Rab5-mCherry). **(K)** Schematic illustrating the bystander BRET assay used to monitor mGα_i_ translocation to subcellular compartments in real time. **(L)** Agonist-dependent translocation of mGα_i_ to subcellular compartments, monitored by real-time bystander BRET in cells co-transfected with FFA4, Nluc-mGα_i_ and Venus-tagged subcellular markers. **(M)** AUC values of mini-G probe translocation to subcellular compartments. Data are mean ± SEM of n = 3 independent experiments.

### FFA4 rapidly internalizes to intracellular compartments in a simple cell model

We next investigated agonist-dependent FFA4 internalization and trafficking in HEK293T cells. As a first approach, we performed live cell imaging experiments in cells transfected with YFP-tagged FFA4 (FFA4-YFP). The cells were imaged by highly inclined thin illumination (HILO) microscopy, which allows for the visualization of dynamic trafficking and signaling events in living cells with high speed and sensitivity^6^. Under basal conditions, FFA4 was localized predominantly at the plasma membrane, with a small fraction at intracellular compartments (Fig. 1D). Upon stimulation with TUG-891, FFA4 rapidly internalized to early endosomes (Fig. 1D), identified by colocalization with Rab5-mCherry (Fig. 1E), and possibly other compartments.

To more precisely characterize and quantify the trafficking profile of FFA4, we then employed a bystander BRET assay that monitors in real-time the proximity between FFA4-Nluc and Venus-tagged subcellular markers^25^ (Fig. 1F). These included K-ras (plasma membrane), Rab5a (early endosomes), Rab4 (fast recycling endosomes), Rab11a/b (slow recycling endosomes), Rab7 (late endosomes/lysosomes), Rab9 (late endosomes to *trans*-Golgi network), Vps29 (retrograde trafficking from early endosome to *trans*-Golgi network), Rab1a (endoplasmic reticulum/ER to the *cis*-Golgi), Rab6 (Golgi and *trans*-Golgi network), and Rab8 (*trans*-Golgi network to the plasma membrane). Basal bystander BRET measurements indicated that, prior to stimulation, FFA4 is present at the plasma membrane (K-ras), recycling endosomes (Rab11a/b), late endosomes (Rab7 and Rab9) and *trans*-Golgi network (Rab8 and Rab9) (Fig. 1G). TUG-891 stimulation caused a rapid redistribution of FFA4 from the plasma membrane (K-ras) and recycling endosomes (Rab11a/b) to early endosomes (Rab5a), late endosomes (Rab7 and Rab9), *trans*-Golgi network (Rab9/Rab6), fast recycling endosomes (Rab4), and the ER (Rab1a). No relevant changes in bystander BRET were detected with retromer (Vps29) or *trans*-Golgi network to plasma membrane (Rab8) markers (Fig. 1H). Overall, similar FFA4 trafficking profiles were obtained with CpdA and α-linolenic acid, even though α-linolenic acid was less effective than CpdA and TUG-891 in inducing FFA4 internalization and subcellular redistribution at the tested 10 μM concentration (Supplementary Fig. 1B).

These results indicated that, in a simple cell system, FFA4 is localized both at the plasma membrane, and to a lesser extent, intracellular compartments under basal conditions and efficiently internalizes and accumulates at intracellular sites after agonist stimulation.

### FFA4 is active at intracellular compartments in a simple cell model

We next asked whether FFA4 is active at intracellular sites in HEK293T cells. To investigate the spatiotemporal pattern of FFA4 activation, we transfected HEK293T cells with FFA4 and Venus-tagged mini-G probes, and imaged them by HILO microscopy. TUG-891 stimulation caused a rapid recruitment of mGα_i_ to the plasma membrane (Fig. 1I), followed by accumulation in early endosomes (Fig. 1J) and other intracellular tubulovesicular structures. To monitor FFA4 activation at specific subcellular compartments, we performed bystander BRET measurements between mini-G probes fused to Nluc and the same panel of Venus-tagged subcellular markers used to follow receptor intracellular trafficking (Fig. 1K). TUG-891 stimulation induced a rapid and strong mGα_i_recruitment to the plasma membrane (K-ras) and, to a variable extent, all investigated intracellular compartments with the exception of the retromer compartment (Vps29) (Fig. 1L, M). CpdA and α-linolenic acid induced overall similar mini-G recruitment profiles, albeit with smaller maximal responses (Supplementary Fig. 1C).

A similar recruitment profile was observed for mGα_o_ (Fig. 1M, Supplementary Fig. 2). In contrast, FFA4 stimulation caused only a modest recruitment of mGα_q_ to the plasma membrane, with negligible recruitment at intracellular compartments (Fig. 1M, Supplementary Fig. 2).

Altogether, these results indicate that, in a simple cell model, FFA4 stimulation with both endogenous and synthetic agonists induces FFA4 activation at the plasma membrane and to a lesser extent various intracellular compartments.

### FFA4 resides at and signals from an intracellular compartment intimately associated with lipid droplets in adipocytes

Having characterized the trafficking and signaling of FFA4 in a simple cell model, we sought to clarify its localization and spatiotemporal signaling pattern in adipocytes. As a first model, we used 3T3-L1 cells, a commonly used pre-adipocyte cell line that can be differentiated in vitro^26^. Since FFA4 is expressed at particularly high levels in brown adipocytes^27^, where lipolysis plays a key role in thermogenesis^28^, we additionally investigated mouse immortalized brown adipocytes that we similarly differentiated in vitro^29^. The endogenous expression of functional FFA4 in differentiated immortalized brown adipocytes was confirmed by the ability of an FFA4 inhibitor/negative allosteric modulator (AH7614) to enhance lipolysis stimulated by the β-adrenergic agonist isoproterenol (Supplementary Fig. 3). The higher transfection efficiency in immortalized brown adipocytes compared to 3T3-L1 cells also allowed us to investigate the coupling specificity of FFA4 by BRET with mini-G probes. The results indicate that, similar to what observed in the simple cell model, FFA4 is predominantly coupled to Gα_i/o_ in differentiated immortalized brown adipocytes (Supplementary Fig. 4).

We next investigated the subcellular localization of YFP-tagged FFA4 expressed in adipocytes. Remarkably, in both differentiated 3T3-L1 and immortalized brown adipocytes, a large proportion of FFA4 was located on intracellular membranes under basal conditions, with only a smaller fraction at the plasma membrane (Fig. 2A, B, Supplementary Fig. 5A). The intracellular compartment containing FFA4 appeared to be intimately associated with lipid droplets, labelled with a fluorescent lipid stain (LipidSpot) (Fig. 2A). Upon stimulation with TUG-891, FFA4 located at the plasma membrane internalized into small endosomal vesicles that were seen trafficking and accumulating near the lipid droplets. Occasionally, vesicles carrying FFA4 appeared to later fuse with the pre-existing intracellular compartment associated with the lipid droplets (Supplementary Fig. 5B).

**Figure 2:**
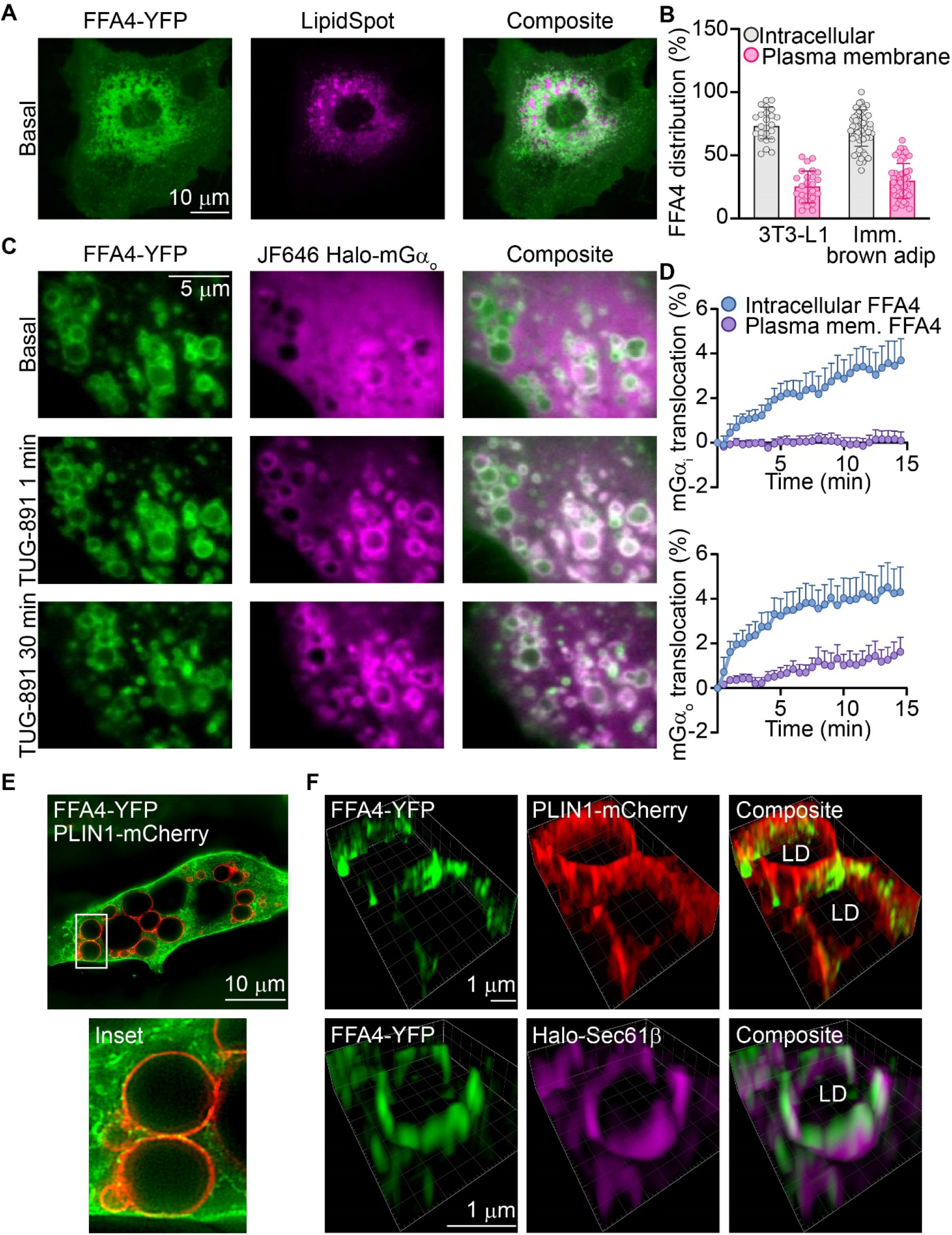
FFA4 resides at and signals from an intracellular compartment intimately associated with lipid droplets in adipocytes. **(A)** Simultaneous imaging of FFA4-YFP with a lipid droplet stain (LipidSpot) in differentiated 3T3-L1 cells. **(B)** Percentage of FFA4-YFP localized at the plasma membrane versus intracellular membranes in differentiated 3T3-L1 and immortalized brown adipocytes. n = 24 and 48 cells for 3T3-L1 and immortalized brown adipocytes, respectively. **(C)** Representative time course of Halo-mGα_o_ recruitment to FFA4-YFP upon stimulation with 10 μM TUG-891 in differentiated 3T3-L1 cells. **(D)** Quantification of mGα_i_ and mGα_o_ translocation to FFA4 at the plasma membrane and intracellular compartment. n = 9 cells for both conditions. **(E)** SIM image of differentiated immortalized brown adipocytes co-transfected with FFA4-YFP and a lipid droplet membrane marker (PLIN1-mCherry). **(F)** Zoomed-in 3D SIM images of differentiated immortalized brown adipocytes co-transfected with FFA4-YFP and PLIN1-mCherry (top) or an ER marker (Halo-Sec61β) (bottom). Data in B and D are mean ± SEM from ≥ 6 independent experiments. LD, lipid droplet.

Remarkably, stimulation with the cell-permeable FFA4 agonist TUG-891 caused a rapid translocation of mGα_i/o_ to the intracellular pool of FFA4 intimately associated with lipid droplets, which for G_o_ occurred as fast as in ∼ 1 min, a timing that precedes FFA4 internalization and appearance in new endosomal vesicles (Fig. 2C, D, and Supplementary Fig. 5A). Of note, mGα_i/o_ translocation was mainly observed to the intracellular FFA4 compartment intimately associated with lipid droplets, with only a minor translocation of mGα_o_ to FFA4 at the plasma membrane and no detectable membrane translocation in the case of mGα_i_.

Altogether, these results indicate that, in adipocytes, a relevant fraction of FFA4 resides in close proximity to lipid droplets prior to stimulation, where it can be rapidly activated by cell-permeable agonists.

### FFA4 colocalizes with an ER maker on membranes surrounding lipid droplets

To further investigate the subcellular localization of FFA4 in adipocytes, we resorted to superresolution imaging by structured illumination microscopy (SIM). Superresolved 3D images obtained in differentiated immortalized brown adipocytes co-transfected with FFA4-YFP and mCherry-tagged perilipin 1 (PLIN1), used as a lipid droplet membrane marker, confirmed that FFA4 is located in close proximity to the surface of lipid droplets (Fig. 2E, F).

Since the lipid droplet membrane consists of a single phospholipid monolayer that is unlikely to accommodate integral membrane proteins like GPCRs, we reasoned that FFA4 should rather be localized on membranes of another, closely associated, membrane compartment. Given the known intimate association of the ER with lipid droplets^30^ and the pattern observed in our SIM images, the ER appeared as a likely candidate. We therefore conducted additional SIM experiments in cells co-transfected with FFA4-YFP and an ER marker (Halo-Sec61β) (Fig. 2F). The results showed a distinct colocalization of FFA4 with Sec61β−positive ER subdomains.

These results suggested that the lipid-droplet associated intracellular FFA4 pool found in adipocytes is likely located on membrane subdomains of the ER.

### Endogenous FFAs released during lipolysis activate FFA4 in adipocytes

Based on our results, we hypothesized that the intracellular FFA4 pool might be rapidly activated by FFAs released following lipolysis induction. To explore this hypothesis, we first compared FFA4 activation with TUG-891 and indirect lipolysis induction with either the β-adrenergic agonist isoproterenol or the adenylyl cyclase activator forskolin, which we measured by BRET with mGα_i/o_ probes. In undifferentiated immortalized brown adipocytes, which only contain few lipid droplets and have negligible lipolytic activity, direct FFA4 activation with TUG-891 caused a rapid and robust increase in BRET, whereas no detectable response was observed upon forskolin or isoproterenol treatment (Fig. 3A). In contrast, in differentiated, lipid droplet-containing immortalized brown adipocytes, both forskolin and isoproterenol stimulation induced a strong and rapid mGα_i/o_ recruitment to FFA4 with a near-maximal response after 2 min, comparable with TUG-891 (Fig. 3A). Consistently, pharmacological stimulation of lipolysis with SR-3420, which binds CGI-58 causing its release from PLIN and subsequent activation of adipose triglyceride lipase (ATGL) independently of cAMP/protein kinase A^31^, also induced mGα_i/o_ but not mGα_s/q/12_ recruitment to FFA4 (Fig. 3B).

**Figure 3:**
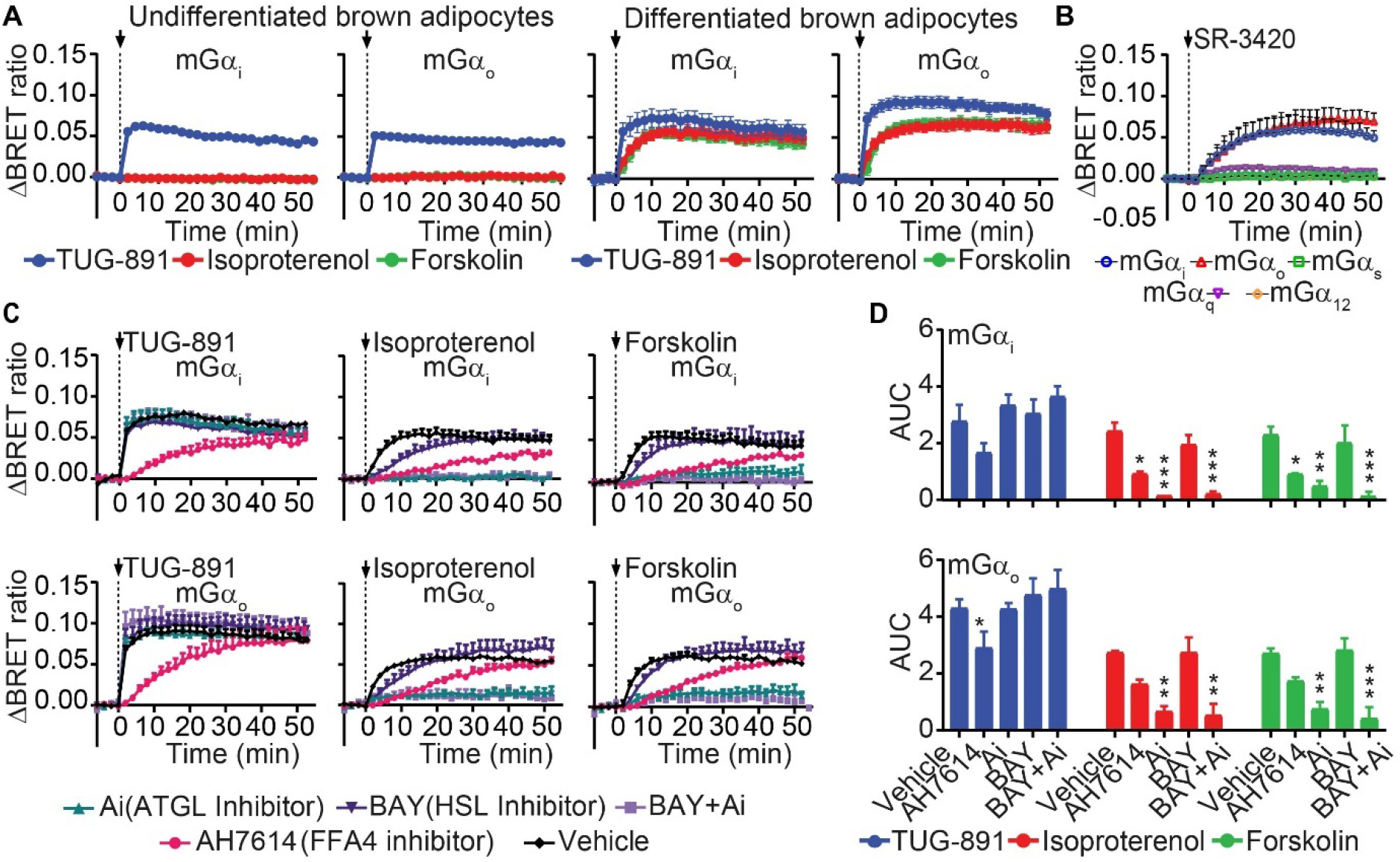
Endogenous FFAs released upon lipolysis activate FFA4 in immortalized brown adipocytes. **(A)** Real-time BRET measurements of FFA4 activation in undifferentiated (left) or differentiated (right) immortalized brown adipocytes. Shown are the results of BRET measurements of Venus-tagged mGα_i/o_ recruitment to FFA4-Nluc upon stimulation with 10 μM TUG-891, isoproterenol or forskolin. **(B)** Venus-tagged mini-G probe recruitment to FFA4-Nluc upon stimulation with 40 μM SR-3420. **(C)** Venus-tagged mini-G probe recruitment to FFA4-Nluc upon stimulation with 10 μM TUG-891, isoproterenol or forskolin after pre-treatment (20 min) with an ATGL inhibitor (Ai, 10 μM), HSL inhibitor (BAY, 5 μM), FFA4 inhibitor (AH7614, 10 μM) or vehicle. **(D)** Corresponding AUC values. Data are mean ± SEM of n = 3 independent experiments. Differences are statistically significant by two-way ANOVA. ****p<0.0001, *** p<0.001, ** p<0.01, * p<0.05 vs vehicle by Du≥nnett’s post hoc test.

To further investigate whether the rapid activation of FFA4 observed in response to forskolin or isoproterenol stimulation is due to lipolysis and the resulting release of FFAs, we took advantage of two lipolysis inhibitors. The first, Atglistatin (Ai), targets ATGL – the enzyme that catalyzes the first and rate-limiting step of triglyceride hydrolysis^32^. The second, BAY 59-9435 (BAY), targets the hormone sensitive lipase (HSL) – catalyzing the second key step of lipolysis^32,33^. Pharmacological inhibition of ATGL with Ai virtually blocked the activation of FFA4 induced by isoproterenol or forskolin (Fig. 3C, D), whereas inhibition of HSL with BAY delayed the response but did not affect its amplitude (Fig. 3C, D). Importantly, neither treatment alone or in combination affected the direct activation of FFA4 with TUG-891, used as a control (Fig. 3C, D). Furthermore, all responses were similarly delayed and reduced by addition of the FFA4 inhibitor AH7614 (Fig. 3C, D).

Altogether, these results indicate that the FFAs released from lipid droplets following lipolysis stimulation in differentiated adipocytes can rapidly induce FFA4 activation and Gα_i/o_ coupling.

### Endogenously released FFAs activate the lipid droplet-associated FFA4 pool to locally modulate cAMP signaling

We then asked whether the rapid activation of FFA4 by endogenously released FFAs occurred via an ‘intracrine’ as opposed to an autocrine/paracrine mechanism and whether FFA4 activation was capable of locally controlling downstream cAMP signaling.

To begin with, we measured FFAs released into the culture medium by a sensitive gas chromatography mass spectrometry (GC-MS) method (detection limit ∼300-400 pM). Under the experimental conditions used for the above BRET experiments, FFA concentrations in the culture medium remained undistinguishable from a blank control sample even after prolonged stimulation with 10 μM forskolin or isoproterenol for 60 min (Fig. 4A, Supplementary Fig. 6). Detectable amounts of palmitoleic (C16:1n7), palmitic (C16:0), oleic (C18:1n-9), vaccenic (C18:1n-7), and myristic (C14:0) acid were only present upon addition of bovine serum albumin (BSA) to the culture medium to facilitate FFA export and accumulation (Fig. 4A, Supplementary Fig. 6)^34,35^. We then measured mGα_o_ recruitment to FFA4 in response to increasing concentrations of palmitoleic, palmitic, oleic, and vaccenic acid, which are reported FFA4 agonists^36-38^. Under the experimental conditions used in our study, detectable FFA4 activation was only observed for FFA concentrations in the micromolar range (Fig. 4B). This is much higher than the upper bound of our estimate of the concentrations of FFAs in the culture medium after forskolin or isoproterenol stimulation. Of note, although palmitic acid was previously reported to be a FFA4 agonist in some^39^ but not all studies^36^, no response was detected for palmitic acid concentrations up to 100 μM, suggesting that it is not a physiologically relevant FFA4 agonist. These results indicate that the very low concentrations of FFAs accumulating in the cell culture medium after lipolysis induction are insufficient to induce any relevant activation of cell-surface FFA4, strongly supporting an intracrine mode of action.

**Figure 4:**
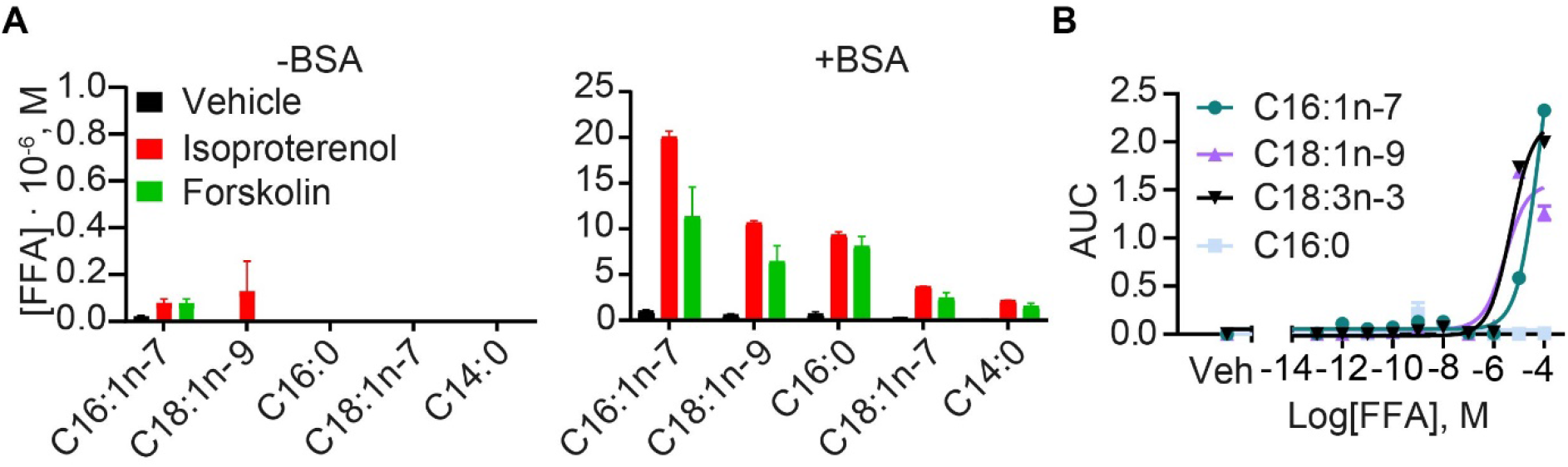
FFA released in the culture medium are insufficient to activate cell-surface FFA4. **(A)** Extracellular fatty acid concentrations detected by GC-MS after stimulation with 10 μM isoproterenol or forskolin for 60 min in the absence (left) or presence (right) of fatty-acid-free BSA (100 μM). **(B)** FFA4 activation in response to increasing concentrations of C16:1n-7 (palmitoleic acid), C18:1n-9 (oleic acid), C18:3n-3 (α-linolenic acid), or C16:0 (palmitic acid) monitored by BRET between FFA4-Nluc and Venus-tagged mGα_o_. Data are mean ± SEM of n = 3 independent experiments.

We therefore attempted to compare the effects of FFA4 on local cAMP signaling at the plasma membrane and intracellular sites. For this purpose, we transfected differentiated immortalized brown adipocytes with a BRET sensor for cAMP, Nluc-Epac-VV^40^, which we tethered to the plasma membrane, ER, or lipid droplet membrane via fusion to targeting domains derived from PDE2A3^41^, Sec61β^42^, or PLIN1^43^, respectively (Fig. 5A). As expected, pharmacological stimulation with either isoproterenol or forskolin induced a rapid reduction of BRET at all compartments (Fig. 5B), indicative of local increases in cAMP levels. Co-transfection of FFA4 attenuated the BRET responses, consistent with FFA4-mediated activation of Gα_i/o_-proteins (Fig. 5B). Remarkably, however, the reduction of cAMP levels caused by the presence of FFA4 was significantly higher at lipid droplets and the ER than at the plasma membrane (lipid droplet > ER > plasma membrane) (Fig. 5B, C). These results indicate that FFA4 exerts a stronger inhibitory effect on cAMP levels near lipid droplets. Of note, FFA4 overexpression had a greater inhibitory effect after stimulation with isoproterenol compared to forskolin, suggesting that FFA4 and β-adrenergic receptors might be preferentially coupled to a shared sub-pool of adenylyl cyclases.

**Figure 5:**
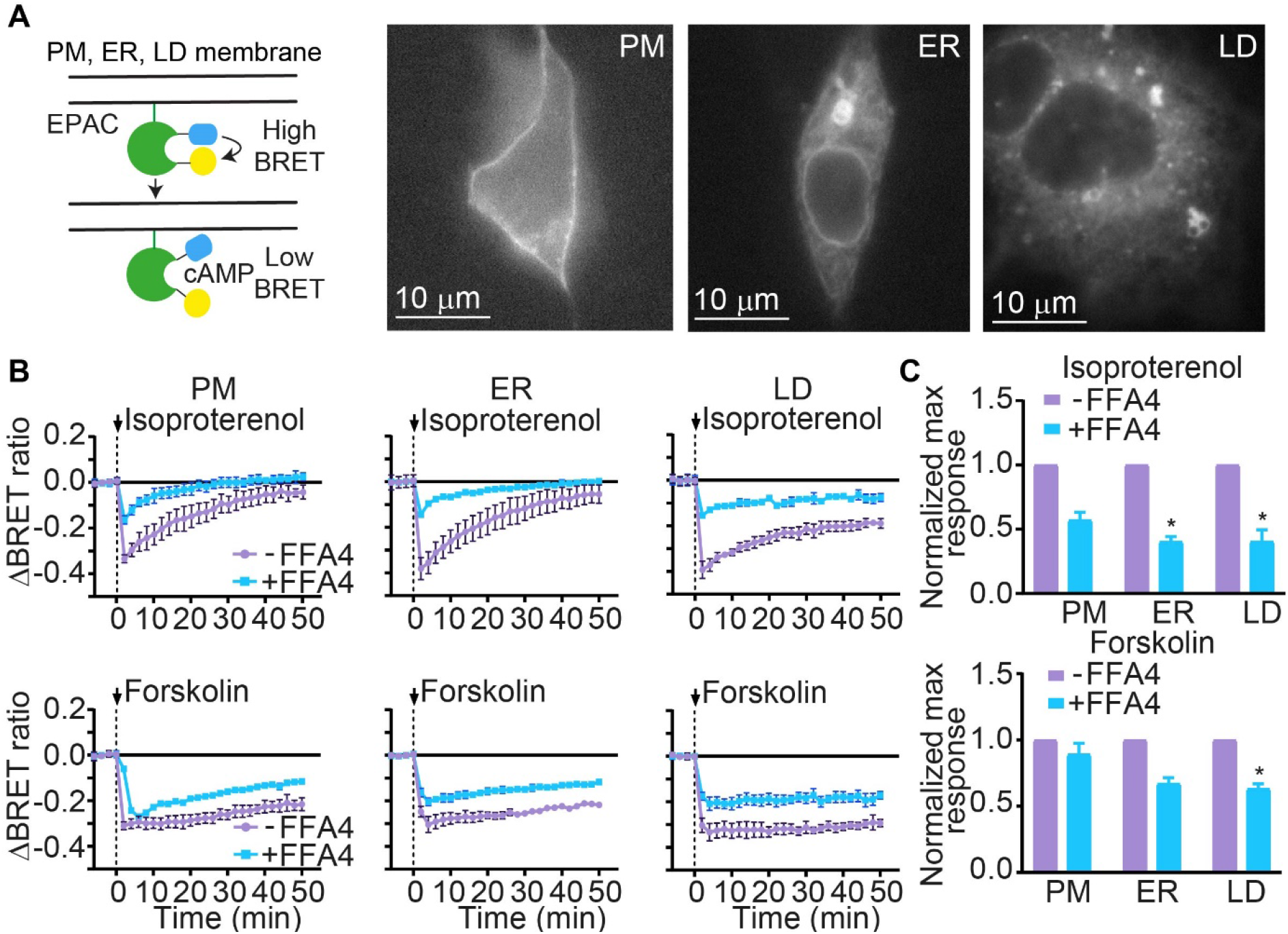
FFA4 exerts a local control over cAMP levels at lipid droplets. **(A)** Left, schematic of the BRET sensor (Nluc-Epac-VV) used to measure changes in local cAMP concentration. Right, representative HILO images of Nluc-Epac-VV tethered to the plasma membrane (PM), endoplasmic reticulum (ER), and lipid droplets (LD). **(B)** Real-time BRET measurements of local cAMP levels in differentiated immortalized brown adipocytes with or without (mock) FFA4 co-transfection, stimulated with 10 μM isoproterenol (top) or forskolin (bottom). **(C)** Quantification of the BRET changes in B. Results are normalized to the corresponding condition without FFA4 co-transfection. Data are mean ± SEM of n = 3 independent experiments. Differences are statistically significant by one-way ANOVA. * p<0.05 vs PM +FFA4 by Dunnett’s multiple comparison post hoc test.

To further test our hypothesis that FFA4 is activated at sites near to lipid droplets upon lipolysis induction, we performed HILO-microscopy experiments in differentiated immortalized brown adipocytes co-transfected with FFA4-YFP and Halo-mGα_o_. Remarkably, lipolysis induction with forskolin or isoproterenol caused a rapid translocation of Halo-mGα_o_ to the intracellular FFA4 pool located in close proximity to lipid droplets, which was detectable in less than 5 min after stimulation (Fig. 6A, B). In contrast, no detectable translocation to the plasma membrane was observed (Fig. 6A, B). A similar response was observed following isoproterenol stimulation in the presence of a dynamin inhibitor (Dyngo 4a), widely used to prevent receptor internalization (Fig. 6C), ruling out that the rapid appearance of active receptors close to lipid droplets was due to internalization and trafficking of FFA4 from the plasma membrane. Likewise, direct lipolysis induction with SR-3420 caused a rapid and robust translocation of Halo-mGα_o_ to the intracellular FFA4 pool (Fig. 6D), further demonstrating that FFAs released by lipolysis activate intracellular FFA4 receptors intimately associated with lipid droplets.

**Figure 6:**
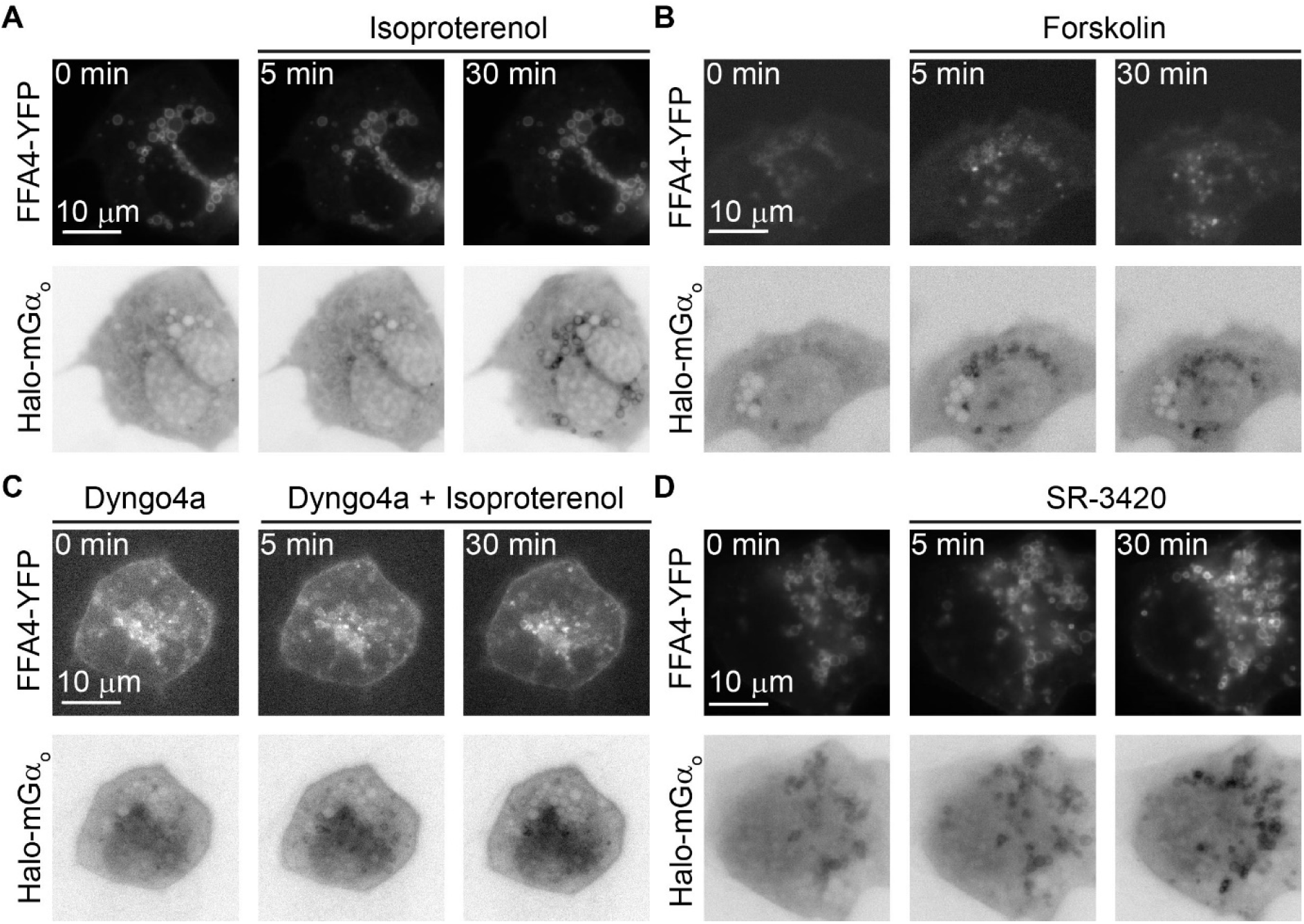
The intracellular pool of FFA4 associated with lipid droplets is rapidly activated upon lipolysis induction in immortalized brown adipocytes. **(A-D)** Representative HILO images of Halo-mGα_o_ recruitment to FFA4-YFP in differentiated immortalized brown adipocytes upon stimulation with 10 μM isoproterenol **(A)**, 10 μM forskolin **(B)**, 10 μM isoproterenol in the presence of 50 μM Dyngo 4a **(C)**, or 40 μM SR-3420 **(D)**. The Halo-mGα_o_ LUT is inverted to facilitate visualization of Halo-mGα_o_ translocation.

Altogether, these findings demonstrate that, upon lipolysis induction in immortalized brown adipocytes, the endogenously released FFAs rapidly bind to and activate an intracellular pool of FFA4 that resides in close proximity to lipid droplets where it can locally couple to Gα_i/o_ and inhibit cAMP signaling, modulating lipolysis in an intracrine fashion.

## DISCUSSION

Our findings on FFA4 in adipocytes reveal a new scenario whereby a metabolite-sensing GPCR is located and signals very close to the intracellular site where its endogenous ligand is produced, allowing it to rapidly modulate a key metabolic pathway in a local fashion (Supplementary Fig. 7). These results not only provide new insights into the mechanisms used by FFA4 to rapidly regulate lipolysis, but also strong evidence for the occurrence of a novel modality of intracrine GPCR signaling by metabolite-sensing GPCRs, a mechanism that has been hypothesized but, to the best of our knowledge, never directly demonstrated.

For more than 40 years, the FFAs released from adipocytes have been known to inhibit lipolysis^44-46^, but the underlying mechanisms have long been elusive. Recently, our team provided evidence that the anti-lipolytic effects of FFAs are at least partially mediated by FFA4^17^. The present study provides novel and unexpected insights into these fundamental mechanisms, which play a key role in physiology and disease and could be exploited for the therapy of metabolic disease.

First, we show that despite being generally classified as a G_q_-coupled receptor^18,19^, FFA4 has a major coupling to Gα_i/o_, which explains its rapid negative effects on lipolysis via inhibition of intracellular cAMP signaling, consistent with our recent observations^17^. While our results highlight the importance of Gα_i/o_ signaling in the control of lipolysis by FFA4, it is worth noting that they do not exclude a contribution of G_q_-coupling to other biological effects of FFA4 in adipocytes and/or other cells. Besides being involved in the FFA4-dependent stimulation of glucose uptake in adipocytes, Gα_q_-coupling is probably particularly relevant in enteroendocrine cells, where it modulates incretin release^36^.

Second, our findings indicate that, in differentiated adipocytes, FFA4 has an unexpected localization on membranes that are intimately associated with lipid droplets, most likely subdomains of the ER. This is consistent with the biogenesis of lipid droplets from the ER and the fact that the two organelles maintain tight functional interactions supported by specialized ER-lipid droplet contact sites^47^. Intriguingly, the unique spatiotemporal organization of FFA4 signaling at lipid droplets appears to be specific to adipocytes, as indicated by our finding that FFA4 is mainly located and signals at the plasma membrane in a simple cell model. Whether this subcellular localization extends to other receptors and/or lipid droplet-containing cells remains to be investigated. Additionally, the apparent shift of FFA4 subcellular localization from the plasma membrane in undifferentiated cells to intracellular sites in differentiated adipocytes may form part of the highly regulated adipocyte maturation program associated with lipid droplet biogenesis. In agreement with this hypothesis, FFA4 expression is strongly induced during adipocyte maturation^48,49^ and has been shown to promote lipid droplet biogenesis^49,50^. Moreover, FFA4 is one of genes that are most strongly upregulated by exposure to cold in brown adipose tissue (BAT)^51,52^. Future studies will be required to investigate the complex interplay between the mechanisms that control FFA4 expression, its subcellular localization and the formation of lipid droplets during adipocyte differentiation.

Third and most importantly, our results show that the endogenous FFAs released during lipolysis can rapidly activate the intracellular pool of FFA4 that is residing on intracellular membranes intimately associated with lipid droplets, thus allowing FFA4 to locally inhibit cAMP production via Gα_i/o_ signaling. The occurrence of G protein signaling at the ER is consistent with the well-known presence of G proteins on ER membranes. In fact, G proteins are assembled and become associated with lipid bilayers on membranes of the ER, allowing them to reach the plasma membrane via anterograde trafficking^53-55^. Similarly, adenylyl cyclases are synthesized in the ER and have been shown to be abundant on its membranes^56^. Intriguingly, recent proteomics have also detected the presence of G proteins, including Gα_s_, Gα_i2_, Gα_q_ and Gβ3 subunits in lipid droplet fractions^57^. Our findings expand these previous observations by showing that, in addition to the whole machinery required for G protein signaling and cAMP production, FFA4 is also present and active in the vicinity of lipid droplets, providing a canonical, receptor-dependent mechanism to locally modulate cAMP signaling at this key intracellular compartment.

Of note, lipid droplets differ substantially between white adipocytes, which are univacuolar, and brown adipocytes, which contain multiple, small lipid droplets. Moreover, the composition of their membrane can vary among cells, lipid droplets within a given cell or even submdomains within a single lipid droplet^58^. This can give rise to distinct subpopulations of lipid droplets within a single cell that co-exist in multiple functional states. It is therefore tempting to speculate that this newly discovered modality of intracrine FFA4 signaling likely evolved to allow adipocytes to rapidly and efficiently fine-tune lipolysis in response to locally released FFAs and, potentially, allow each lipid droplet to independently sense the FFAs released from its own store and adjust its lipolytic flux accordingly.

Given the potential of FFA4 as a drug target for the treatment of metabolic and inflammatory disease^24,59,60^, our findings could also have important therapeutic implications. Above all, they indicate that drugs exploiting FFA4 antilipolytic effects should cross the plasma membrane and access the intracellular FFA4 pool associated with lipid droplets to efficiently modulate lipolysis. Moreover, the fact that FFA4 appears to signal from multiple subcellular compartments that differ among cell types opens up the possibility of selectively activating or inhibiting FFA4 signaling at specific subcellular locations to induce more specific pharmacological responses while minimizing unwanted effects.

Altogether, our study reveals the occurrence of a previously unrecognized intracrine signaling modality by a prototypical metabolite-sensing GPCR and the existence of a specialized intracellular signaling hub that is intimately associated with lipid droplets and provides a platform to rapidly and efficiently regulate lipolysis in response to local FFA release.

## METHODS

### Plasmids

A plasmid containing FFA4-eYFP was as previously described^19^. The FFA4 sequence was subcloned into pcDNA3 following BamHI and EcoRI digestion. Nluc was subcloned from mG-Nluc using EcoRI and XhoI restriction sites and ligated into the pcDNA3 FFA4 construct to generate FFA4 containing Nluc fused to its C-terminus (FFA4-Nluc). Plasmids encoding mini-G probes were kindly provided by Nevin Lambert, Augusta University^23^, and the PLIN1 construct by David Savage, University of Cambridge^61^. A plasmid encoding Halo-Sec61β was a kind gift from Christopher Obara, Janelia Research Campus^42^ and Rab5-mCherry were kindly provided by Tom Kirchhausen (Harvard Medical School, USA). Plasmids expressing Venus-tagged subcellular markers were kindly provided by Kevin Pfleger, University of Western Australia^25^, and a plasmid encoding the Nluc-EPAC-VV BRET sensor was kindly provided by Kirill Martemyanov, Scripps Research Institute^40^. A plasmid encoding Nluc-Epac-VV with Sec61β^42^ fused to its C-terminus was cloned by PCR and Gibson assembly using Nluc-Epac-VV as a template. Likewise, plasmids encoding Nluc-Epac-VV carrying a PDE2A3 plasma membrane targeting domain^41^ or full length PLIN1^43^ at the N-terminus were cloned by PCR and Gibson assembly using Nluc-Epac-VV as a template.

### Cell culture and transfection

HEK293T cells (ATCC) were cultured in Dulbecco’s Modified Eagle Medium (DMEM) supplemented with 10% fetal bovine serum (FBS), 100 U/ml penicillin and 0.1 mg/ml streptomycin at 37 °C, 5% CO_2_. For BRET experiments, HEK293T cells were plated onto 6-well plates at a density of 7 × 10^5^ cells/well. The next day, cells were transfected with Lipofectamine 2000, following the manufacturer’s protocol. After 24 hours, cells were resuspended in FluoroBrite phenol red-free DMEM medium supplemented with 4 mM L-glutamine and 5% FBS and plated onto poly-D-lysine-coated 96-well white polystyrene Nunc microplates at a density of 1 × 10^5^ cells/well and allowed to adhere for 24 hours. For live-cell imaging assays, HEK293T cells were seeded onto 25-mm round glass coverslips at a density of 5 × 10^5^ cells/well. The next day, cells were transfected with Lipofectamine 2000, following the manufacturer’s protocol. Cells were imaged 24 hours after transfection.

Immortalized mouse brown preadipocytes^29,62^ were cultured in high-glucose GlutaMAX DMEM supplemented with 10% FBS, 100 U/ml penicillin and 0.1 mg/ml streptomycin at 37 °C, 5% CO_2_. Preadipocytes were seeded onto 10-cm dishes at a density of 1.3 × 10^6^ cells/dish, 96-well plates at a density of 6 × 10^4^ cells/well or 25-mm round glass coverslips at a density of 5 × 10^5^ cells/well for BRET measurements, glycerol quantification assays and imaging experiments, respectively. After 48-72 hours, the medium was replaced with complete high-glucose GlutaMAX DMEM containing 500 μM 3-isobutyl-1-methylxanthine (IBMX), 1 μM dexamethasome, 1 nM triiodo-L-thyronine (T3), 0.5 μM rosiglitazone, and 20 nM human insulin for 48 hours, followed by complete high-glucose GlutaMAX DMEM containing 1 nM T3 and 20 nM human insulin for 24 hours to induce differentiation. The following day, differentiated adipocytes were transfected using TransIT-X2 transfection reagent (Mirus) as per the manufacturer’s protocol. For BRET assays, differentiated immortalized brown adipocytes grown on 10-cm dishes were detached with 0.25% trypsin-EDTA, reverse transfected and replated onto Nunc microplates at a density of 7.5 × 10^4^ cells/well in high-glucose GlutaMAX DMEM supplemented with 10% FBS; whilst for imaging experiments, differentiated immortalized brown adipocytes grown on 25-mm round glass coverslips were forward transfected. All transfected adipocytes were incubated for 24 hours before being used for BRET assays or imaging.

3T3-L1 cells (ATCC) were cultured in DMEM supplemented with 10% bovine calf serum, 100 U/ml penicillin and 0.1 mg/ml streptomycin at 37 °C, 5% CO_2_. Preadipocytes were seeded onto 10-cm dishes, and, once confluent, transfected by electroporation. Confluent cells were detached with 0.25% trypsin-EDTA, pelleted and resuspended in 240 μl Dulbecco’s phosphate-buffered saline (DPBS), followed by addition of 40 μg DNA into 4-mm cuvettes. Cells were electroporated at 320 V and 125 μF using a Gene Pulser XCell Eukaryotic system (Bio-Rad), and resuspended in 1 ml of DMEM supplemented with 10% bovine calf serum, 100 U/ml penicillin and 0.1 mg/ml streptomycin. Subsequently, 100 μl of the suspension containing transfected cells were plated onto 25-mm round glass coverslips and allowed to adhere. To induce differentiation, the medium was replaced 48 hours later with DMEM supplemented with 10% FBS, 1 μM dexamethasone, 0.5 mM IBMX, 1.0 μg/ml bovine insulin, 100 U/ml penicillin, and 0.1 mg/ml streptomycin and left for 48 hours. The medium was replaced every 48 hours until cells reached the desired differentiation (7-15 days).

### BRET assays

On the day of the experiment, the medium was replaced with Hanks’ Balanced Salt solution (HBSS) containing 10 mM HEPES pH 7.5 and 10 μM furimazine/Nano-Glo Luciferase Assay Substrate. BRET measurements were performed at 37 °C using a PHERAstar Microplate Reader (BMG Labtech) with a dual-luminescence BRET1 plus readout filter (460-490 nm band-pass, 520-550 nm long-pass). Following 4 baseline measurements, the cells were treated with vehicle or the indicated agonist concentration and measured for an additional hour. BRET acceptor/donor ratios were calculated separately for each well. Shown are the changes in BRET ratio over basal (ΔBRET ratio) after vehicle subtraction. Measurements were performed using at least two technical replicates.

### Live cell protein labeling

Cells were labelled with 1 μM HaloTag Janelia Fluor 646 (JF646) (Promega) in complete culture medium (without antibiotics) for 20 min at 37 °C. Cells were then washed three times with complete culture medium, allowing 5 min incubation between washes.

### HILO live-cell imaging

25-mm round glass coverslips were mounted in a microscopy chamber filled with HBSS supplemented with 10 mM HEPES, pH 7.5. The sample and objective were maintained at 37 °C using a temperature-controlled enclosure throughout the experiments. Live-cell imaging were performed using total internal reflection fluorescence (TIRF) illumination on a custom system (assembled by CAIRN Research) based on an Eclipse Ti2 microscope (Nikon, Japan) equipped with four EMCCD cameras (iXon Ultra 897, Andor), 100x oil-immersion objective (SR HP APO TIRF NA 1.49, Nikon), an iLas2 TIRF illuminator (Gataca Systems), 405, 488, 561, and 637 nm diode lasers (Coherent, Obis), a quadruple beam splitter, quadruple band excitation and dichroic filters, 1.5x tube lens and hardware focus stabilization. Two of the four synchronized EMCCSs were used to acquire simultaneous image sequences at a rate of one image every 30 seconds.

### Structured illumination microscopy (SIM)

Differentiated immortalized brown adipocyte cells were seeded and transfected on precision cover glasses thickness No. 1.5H (Marienfeld). Cells were washed twice with DPBS for 5 min each and fixed with 4% paraformaldehyde in 0.1 M PIPES, pH 6.95, 2 mM EGTA, 1 mM MgSO_4_ for 15 min at room temperature, and quenched with 50 mM ammonium chloride. After fixation, coverslips were air dried before mounting with ProLong Diamond Antifade mountant (Vector Laboratories). Slides were cured at room temperature for 48 hours before imaging. Images were acquired using 2D and 3D SIM and reconstructed in NIS-Elements (Nikon) or ARIVIS (Zeiss), respectively.

### Image-based quantification of mini-G probe translocation

Ilastik (version 1.4.0)^63^ was used to train a random forest to classify pixels from the FFA4 channel into three classes: FFA4, intracellular background and extracellular background. One classifier/class was trained for each cell line, i.e. 3T3-L1 and immortalized brown adipocytes, and all default image features were used. Training images were selected across conditions and replicates.

The remaining analysis was performed using a Fiji (version 2.9.0/1.53t) macro^64^, which is available at https://github.com/JeremyPike/miniG-analysis. First, the probabilities from the ilastik classifier were thresholded at 0.5 for the FFA4 and intracellular background classes to give binary segmentations. FFA4 and intracellular background segmentations were then combined (logical OR) to get a binary cellular segmentation. This cellular segmentation was post processed by removing connected components smaller than 5,000 pixels, morphological closing (circular structural element with 3 pixel radius) and hole filling. The ‘plasma membrane’ region was defined as a 10 pixel deep band at the edge of the cellular segmentation. The percentage of total cellular FFA4 signal (all pixels outside the binary FFA4 segmentation are zeroed) within the membrane region was then quantified for each time point. To quantify mini-G translocation to FFA4 at the plasma membrane and intracellular compartments, we first subtracted the mean extracellular background intensity measured at the first time point from the raw mini-G signal. We then quantified the percentage of total cellular mini-G signal colocalizing with membrane FFA4 (overlap with both the binary FFA4 and cellular segmentations) or intracellular FFA4 (further than 10 pixels from the cellular segmentation boundary). Shown are the changes over basal following TUG-891 stimulation.

### Glycerol quantification

Glycerol was quantified using the luminescence Glycerol-Glo assay (J3151, Promega), as per manufacturer’s protocol. In brief, differentiated immortalized brown adipocytes were serum starved for 2 hours, washed with DPBS, and treated as indicated in Krebs Ringer buffer pH 7.0 with or without 100 μM fatty-acid-free BSA. 50 μl of the cell supernatant was collected at the specified time points and combined with the Glycerol Detection Reagent at a 1:1 ratio, resulting in a luminescent signal that is proportional to the amount of glycerol present. Luminescence was measured using the PHERAstar Microplate Reader (BMG Labtech). The glycerol concentration of each sample was calculated using the luminescence of a glycerol standard with a known concentration.

### Fatty acid extraction

Differentiated adipocytes were serum starved for 2 hours, washed with DPBS, and treated as indicated in Krebs Ringer buffer (pH 7.0) with or without 100 μM fatty-acid-free BSA. Cell supernatants were collected at the specified time points and transferred to pre-chilled glass vials alongside thawed cell-free buffer blanks. Lipids were extracted using a 1:1:2 v/v ratio of methanol (Biosolve) containing 0.01% w/v butylated hydroxytoluene (Sigma), water (Biosolve) and dichloromethane (Sigma) containing heptadecanoate acid (C17:0, Sigma) as an internal standard. The extraction mixture was carefully vortexed followed by phase separation via centrifugation at 11,000 x g for 15 min at 4 ºC. The bottom dichloromethane non-polar lipid-containing layer was transferred to a fresh glass vial using a glass pipette and subsequently dried under a gentle N_2_ stream at room temperature.

### Gas chromatography mass spectrometry (GC-MS)

Non-esterified fatty acids were analyzed following derivatization to their pentaflurobromobenzene esters. Unlike derivatization to fatty acid methyl esters (FAME) this technique does not disrupt esterified fatty acids, eliminating the requirement for pre-analytical separation of these species^65,66^. Dried samples were first resuspended in a 1:1 v/v mixture of a 1% solution of N,N-diisopropylethylamine (Sigma) and a 1% solution of pentaflurobenzylbromine (Sigma) both in acetonitrile (Hypergrade, Supelco)and left at room temperature for 30 min whilst protected from light. The solution was dried under a N_2_ stream and resuspended in 100 μl GC-MS-grade isooctane (SupraSolv, Supelco), vortexed, and transferred to glass vials for GC-MS analysis. A set of analytical standards were created in parallel, whereby commercial saturated/monounsaturated fatty acid (Cayman) and a polyunsaturated fatty acid mixture (Cayman) were prepared in a 1:1 ratio and analyzed within the same analytical run. Pooled quality control samples were also prepared to run at the start, middle and end of each batch.

Samples were analyzed using an Agilent 8890 GC connected to a 5977B MSD system with a G3393B CI Upgrade kit, whereby 5 μl of sample were injected in splitless mode onto a 90-m FastFAME Column (Agilent) using ultrapure helium (BOC) at a constant pressure of 46 psi. The following non-linear temperature gradient was used: 50 °C before ramping to 160 °C at 50 °C /min followed by ramping to 230 °C at 10 °C /min, held for 30 min, before ramping to 250 °C at 1 °C /min, held for 16 min. The MS detector operated in negative chemical ionization mode with ultrapure methane (N5.5, BOC) as the collision gas. Collision gas flow rates and electron energies were determined by an instrument autotune at the start of each analytical run. The transfer line was maintained at 250 °C, MS source at 280 °C and MS quad at 200 °C. Fatty acids were detected as their [M-H]-adducts following electron capture dissociation in full scan mode between 150 and 400 m/z. The lower limits of quantification for each fatty acid species were: C16:1n-7 = 373 pM, C18:1n-9/n-7 = 359 pM, C16:0 = 390 pM, C14:0 = 438 pM.

### Mass spectrometry data processing

Raw MS Spectra were converted to .CDF files using Agilent ChemStation and processed using El Maven (Elucidata). Fatty acids were identified using a combination of observed m/z and retention time matching to both an internal library of authentic lipid standards and commercial standards run as part of the same analytical batch. Peak areas for each fatty acid species were normalized to the peak area of the internal standard Heptadecanoate acid (C17:0) then baseline subtracted against the appropriate buffer blank from the same analytical run. For absolute quantification, calibration lines were constructed by linear regression (GraphPad) before conversion of normalized signal intensity to concentration using either direct comparison (if the fatty acid species was contained within the quantitative standard) or the nearest structurally similar lipid (e.g. for C18:1n-7 the calibration line for C18:1n-9 was used).

### Statistics

Statistical analysis was performed using GraphPad Prism 9 software. Values are given as mean ± SEM. Differences between two groups were assessed by a two-tailed student’s t-test. Differences between three or more groups were assessed by one-way or two-way analysis of variance ANOVA test followed by Dunnett’s post hoc test for multiple comparisons between groups, as appropriate. Differences were considered significant for P values < 0.05.

## ACKNOWLEDGEMENTS

This study was supported by a Wellcome Trust Senior Research Fellowship (212313/Z/18/Z to D.C.). E.T., S.L.O’B., T.M., and D.C. participate in the European COST Action CA18133 (ERNEST). We wish to acknowledge the support and resources of the Birmingham Metabolic Tracer Analysis Core. Mass Spectrometry work was supported by funding from the Cancer Research UK Birmingham Centre award C17422/A25154. AB was supported by Cancer Research UK [SEBSTF-2021\100002].

## AUTHOR CONTRIBUTION

D.C. and S.L.O’B conceived the study. E.T., S.L.O’B., and G.S. performed the BRET experiments and analyzed the data. E.T. performed HILO and SIM imaging experiments. J.C. provided support with HILO and SIM imaging experiments. T.M. assisted with cloning and design of experiments. J.P. developed the computational analyses to quantify HILO images. J.R., and A.B. performed GC-MS experiments, supervised by D.A.T. G.M. and B.D.H. provided reagents and contributed to the design and interpretation of the experiments with pharmacological FFA4 stimulation. Z.G-H., and T.W.S. contributed to the design and interpretation of the experiments in immortalized brown adipocytes.

