## Supplementary data for "A new paradigm of intracrine free fatty acid receptor 4 signaling at lipid droplets"

**SUPPLEMENTARY INFORMATION**

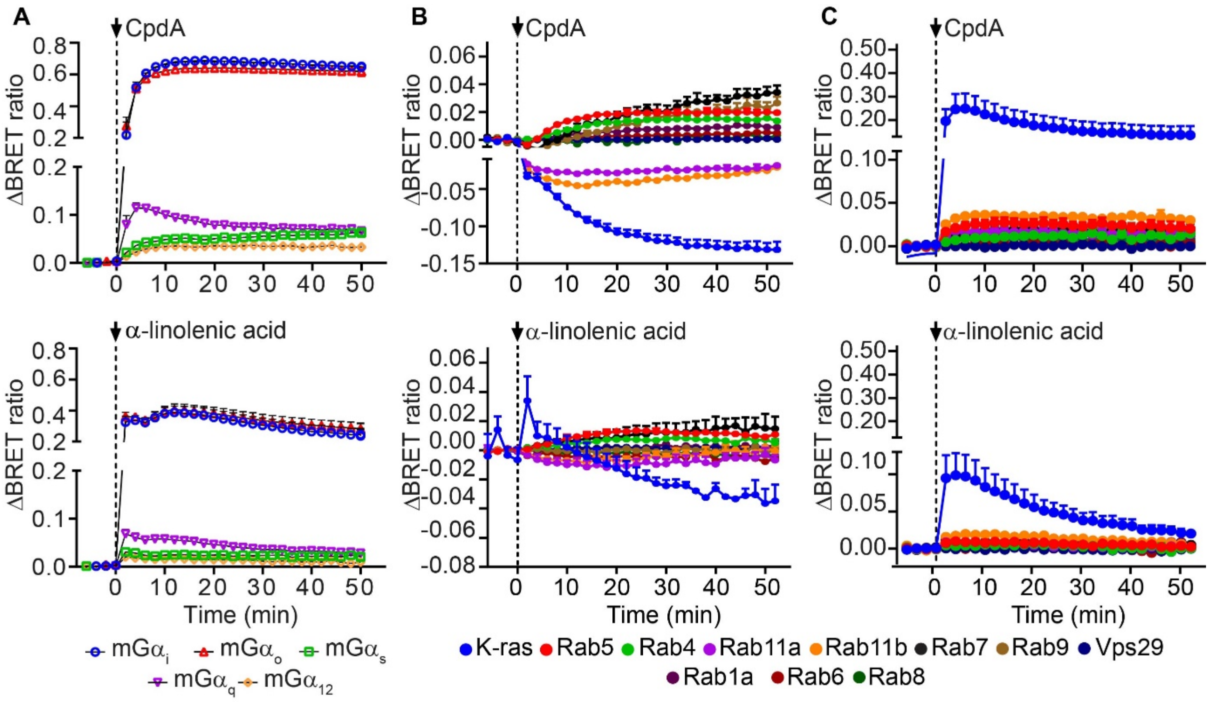


**Supplementary Figure 1: Effects of CpdA and α-linolenic acid stimulation on FFA4 signaling and trafficking.** Shown are the results of real-time BRET measurements in HEK293T cells transfected with the indicated constructs and stimulated with 10 μM CpdA (top) or α-linolenic acid (bottom). **(A)** Agonist-dependent increases in Venus-tagged mini-G probe recruitment to FFA4-Nluc. **(B)** Agonist-dependent changes in FFA4 localization at subcellular compartments, monitored by BRET between FFA4-Nluc and Venus-tagged subcellular markers. **(C)** Agonist-dependent translocation of mG$\alpha_i$ to subcellular compartments in cells co-transfected with FFA4, Nluc-mG$\alpha_i$ and Venus-tagged subcellular markers. Data are mean ± SEM of n = 3 independent experiments.

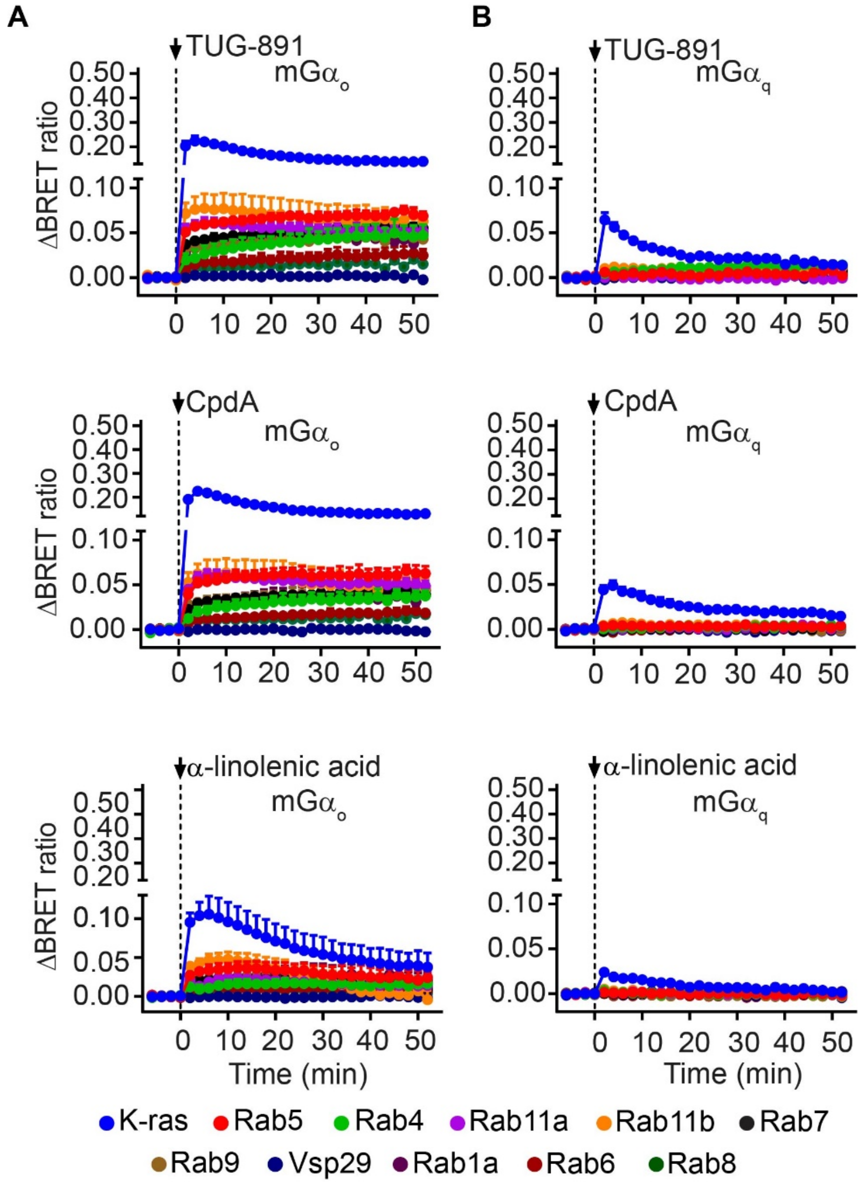


**Supplementary Figure 2: $mG\alpha_o$ and $mG\alpha_q$ recruitment to intracellular compartments upon FFA4 activation in a simple cell model. (A, B)** Agonist-dependent changes in $mG\alpha_o$ **(A)** or $mG\alpha_q$ **(B)** translocation to subcellular compartments upon stimulation with 10 μM TUG-891 (top), CpdA (middle), α-linolenic acid (bottom), monitored by real-time BRET in HEK293T cells co-transfected with FFA4, Nluc-mGα and Venus-tagged subcellular markers. Data are mean ± SEM of n = 3 independent experiments.

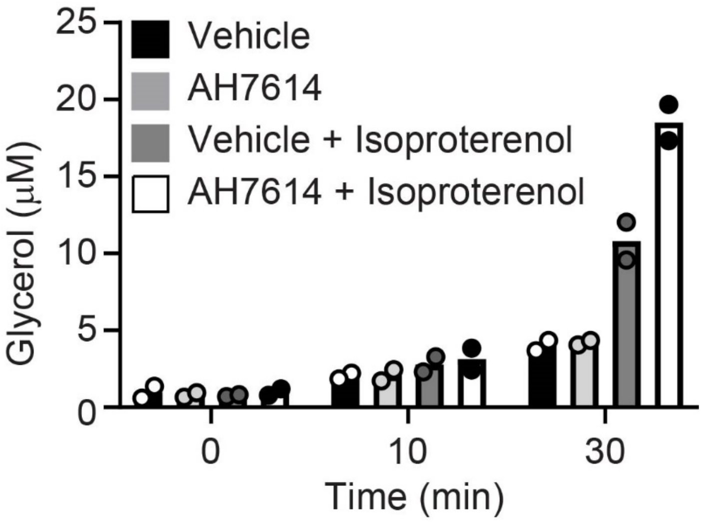


**Supplementary Figure 3: Endogenous FFA4 inhibits lipolysis in differentiated immortalized brown adipocytes.** Differentiated immortalized brown adipocytes were stimulated with 500 pM isoproterenol with or without 15 min preincubation with a FFA4 inhibitor (AH7614, 10 μM). Lipolysis was assessed by measuring glycerol release into the cell culture medium. n = 2 independent experiments.

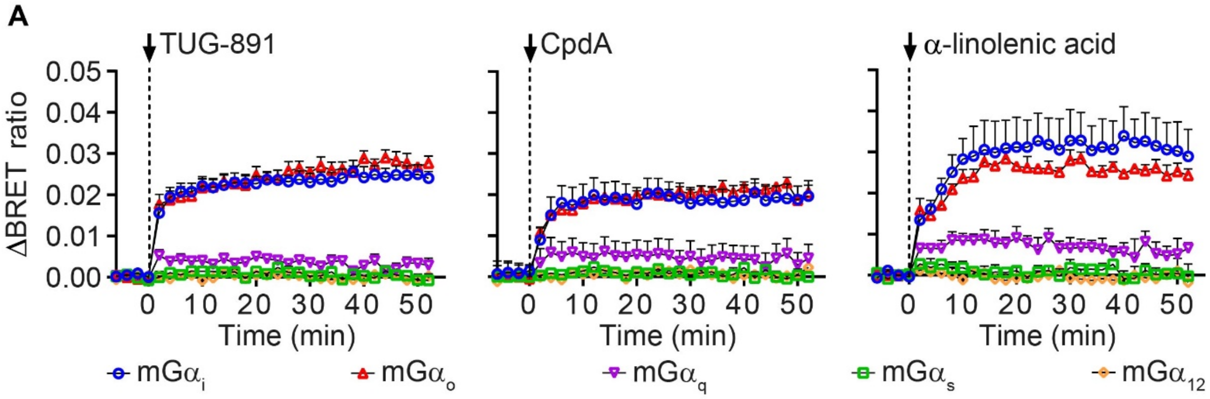


**Supplementary Figure 4: FFA4 is predominantly coupled to $G\alpha_{i/o}$ in differentiated immortalized brown adipocytes. (A)** Agonist-dependent increases in Venus-tagged mini-G probe recruitment to FFA4-Nluc upon 10 μM TUG-891 (left), CpdA (middle), α-linolenic acid (right) stimulation, monitored by real-time BRET measurements in differentiated immortalized brown adipocytes cells. Data are mean ± SEM of n = 3 independent experiments.

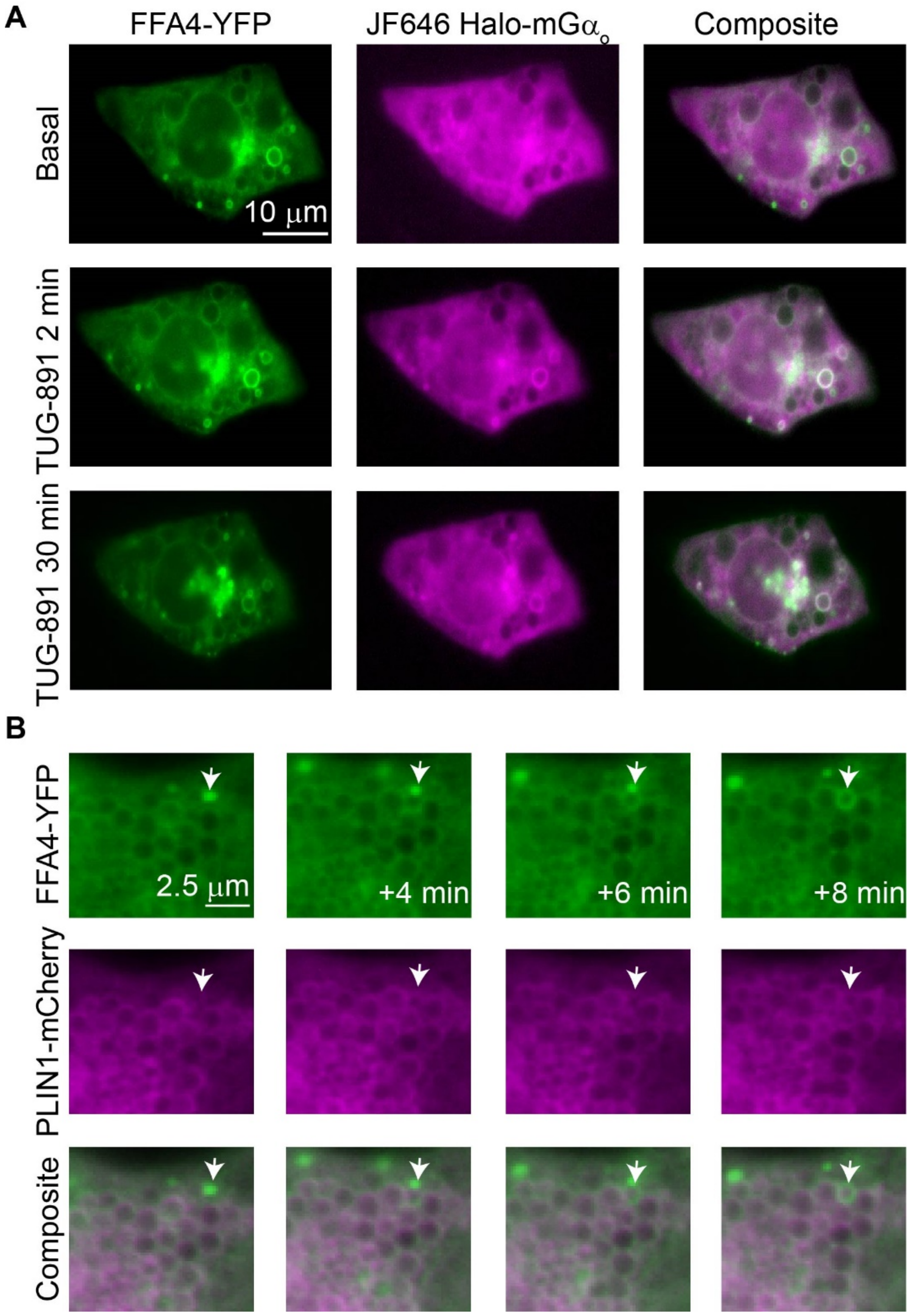


**Supplementary Figure 5: FFA4 localization and signaling in differentiated immortalized brown adipocytes. (A)** Representative time course HILO images of Halo-mGα$_o$ recruitment to FFA4-YFP upon stimulation with 10 µM TUG-891 in differentiated immortalized brown adipocytes. **(B)** Representative HILO image sequence showing a vesicle carrying FFA4-YFP apparently fusing with membranes belonging to the lipid-droplet associated compartment (arrow) in a differentiated immortalized brown adipocyte stimulated with 10 µM TUG-891.

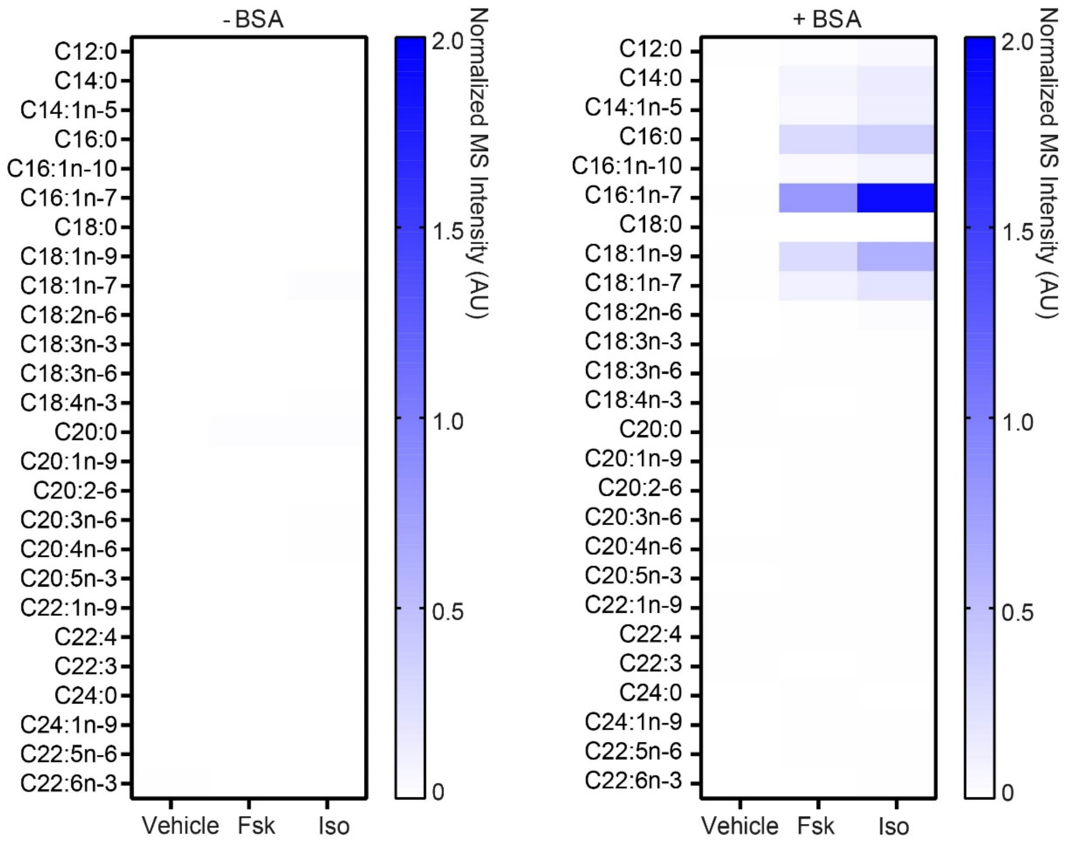


**Supplementary Figure 6: Fatty acid species released by differentiated immortalized brown adipocytes.** Shown are heat maps reporting the normalized GC-MS intensities of the indicated FFAs measured in the culture medium after stimulation with 10 µM forskolin (Fsk) or isoproterenol (Iso) for 60 min in the absence (left) or presence (right) of 100 µM fatty-acid-free BSA.

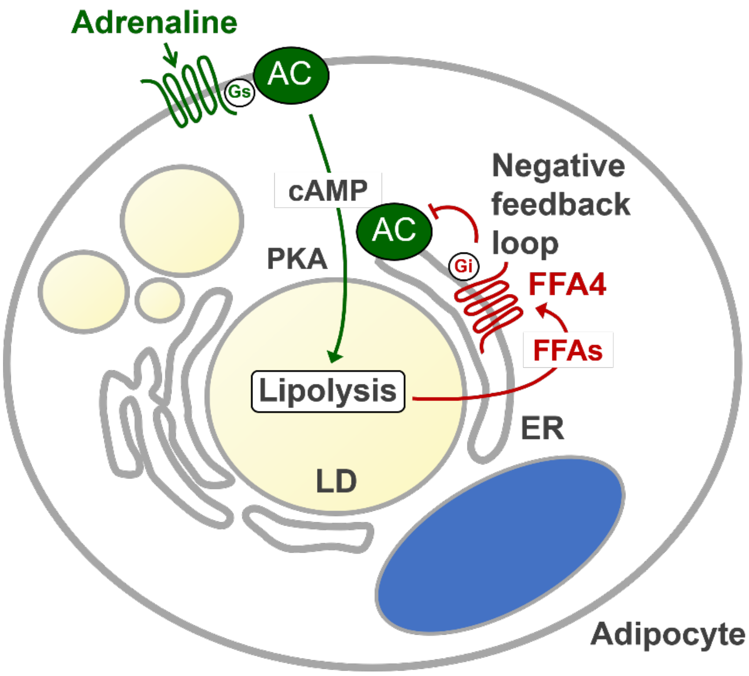


**Supplementary Figure 7: Cartoon summarizing the new paradigm of 'intracrine' FFA4 signaling at lipid-droplets as revealed by the present study.** Adrenaline, like other hormones that signal through Gs coupled-receptors, activates the lipolytic machinery located at the surface of a lipid droplet via stimulation of the cAMP/PKA signaling pathway, with consequent local release of FFAs. The released FFAs rapidly bind to an intracellular pool of FFA4 that resides on membranes of the endoplasmic reticulum (ER) that are intimately associated with the lipid droplet. FFA4, in turn, inhibits via Gi/o coupling the production of cAMP near the surface of the lipid droplet, ultimately reducing its lipolytic rate. This provides a previously unknown local feedback mechanism that allows each lipid droplet to rapidly and precisely sense the FFAs released from its own store and adjust its metabolic flux accordingly.
